# Modular automated microfluidic cell culture platform reduces glycolytic stress in cerebral cortex organoids

**DOI:** 10.1101/2022.07.13.499938

**Authors:** Spencer T. Seiler, Gary L. Mantalas, John Selberg, Sergio Cordero, Sebastian Torres-Montoya, Pierre V. Baudin, Victoria T. Ly, Finn Amend, Liam Tran, Ryan N. Hoffman, Marco Rolandi, Richard E. Green, David Haussler, Sofie R. Salama, Mircea Teodorescu

## Abstract

Organ-on-a-chip systems combine microfluidics, cell biology, and tissue engineering to culture 3D organ-specific *in vitro* models that recapitulate the biology and physiology of their *in vivo* counterparts. Here, we have developed a multiplex platform that automates the culture of individual organoids in isolated microenvironments at user-defined media flow rates. Programmable workflows allow the use of multiple reagent reservoirs that may be applied to direct differentiation, study temporal variables, and grow cultures long term. Novel techniques in polydimethylsiloxane (PDMS) chip fabrication are described here that enable features on the upper and lower planes of a single PDMS substrate. RNA sequencing (RNA-seq) analysis of automated cerebral cortex organoid cultures shows benefits in reducing glycolytic and endoplasmic reticulum stress compared to conventional *in vitro* cell cultures.

## Introduction

Cell culture has been a principal model for studying human disease and development for over 70 years since the isolation of HeLa cells from a human cervical cancer biopsy^1,2^. Originally used as a vehicle to research viruses, human cell culture protocols were optimized for quick and easy growth to produce large quantities of material. While tissue culture protocols have advanced, particularly in reducing many media components, much of these original recipes remain in place. There is still much room to improve for tissue culture protocols to mimic physiological nutrient concentrations, supply, and removal. Automated microfluidics allows us to move beyond traditional protocols by feeding at rates and precision not manually feasible.

Recent advances in stem cell and developmental biology have generated more accurate models resembling aspects of primary human tissue^3,4^. Human embryonic stem (hES) cells and induced pluripotent stem (iPS) cells, collectively referred to as pluripotent stem cells (PSCs), have the potential to differentiate into most cell types of the body, and protocols capitalized upon this to generate 3D culture models for human tissues including brain, gut, liver and breast^5,6^. These self-assembling, organ-specific cell cultures, called organoids, are broadly utilized as *in vitro* models in developmental research, pathogenesis, and medicine^7–9^. Organoids mimic key functional characteristics of their primary tissue counterpart’s physiology more accurately than 2D cell cultures^6^. Beyond use as *in vitro* models, organoids are also being explored for applications in regenerative medicine, and healthcare as tissue for implants^5^. As the technology opens up new frontiers, there is a raising need for better methods to grow, control, and analyze organoid cultures. There is a need to lessen the gap between *in vitro* cultures and primary tissue. Efforts like the advances shown here leverage precision microfluidics, robotic automation, and contact-less sensing to enable robust, reproducible cell culture optimized for tissue fidelity.

The ability of PSC-derived organoids to self-assemble and generate many tissue-specific cell types makes them particularly useful for modeling complex tissues and systems. The brain contains some of the highest complexity in the human body, and researchers can generate high-quality models of different brain regions with organoid technology. Cerebral organoids are a form of brain organoid that models the physiology of the cerebral cortex, containing many cortex-specific cell types and sub-regions^10^. These organoids are widely used for research on prenatal brain development^11–13^, brain pathologies^14^, and therapeutic testing^15^. Cerebral organoids will grow to a few millimeters in diameter during prolonged culture and can be maintained in culture indefinitely^16^.

Figure 1 illustrates the process of generating human cerebral organoids by aggregating human PSCs^17,18^. Neural induction is achieved by inhibiting the WNT (IWR1-ε) and Nodal/Activin (SB431542) pathways that yield both dorsal and ventral cortical tissue. As organoid development proceeds at a pace similar to primary fetal development, these cultures must be maintained for weeks to months to observe late-stage cell types and tissue structures. These include terminally differentiated cell types such as neurons, astrocytes, and oligodendrocytes, and the observation of locally-organized rosettes comprised of radial glia neural stem cells (Figure 1B). Current protocols of cerebral organoids have limitations in throughput, consistency, and reliability, as well as impaired health, showing hallmarks signs of cellular stress^19,20^. The phenotype of cell stress is likely multifactorial; however, many factors may result from traditional cell culture. The pace of manual media changes (often once every 1-3 days) leads to erratic swings in nutrient availability and the build-up of toxic metabolites. By feeding outside the incubator, cells experience colder temperatures and reduced dissolved carbon dioxide leading to a more alkaline environment. Lab automation is becoming increasingly prevalent in life sciences to reduce the effect of these uncontrolled variables and increase the quantity and quality of experiments enabling easy long-term maintenance with minimal manual requirements.

**Figure 1.**
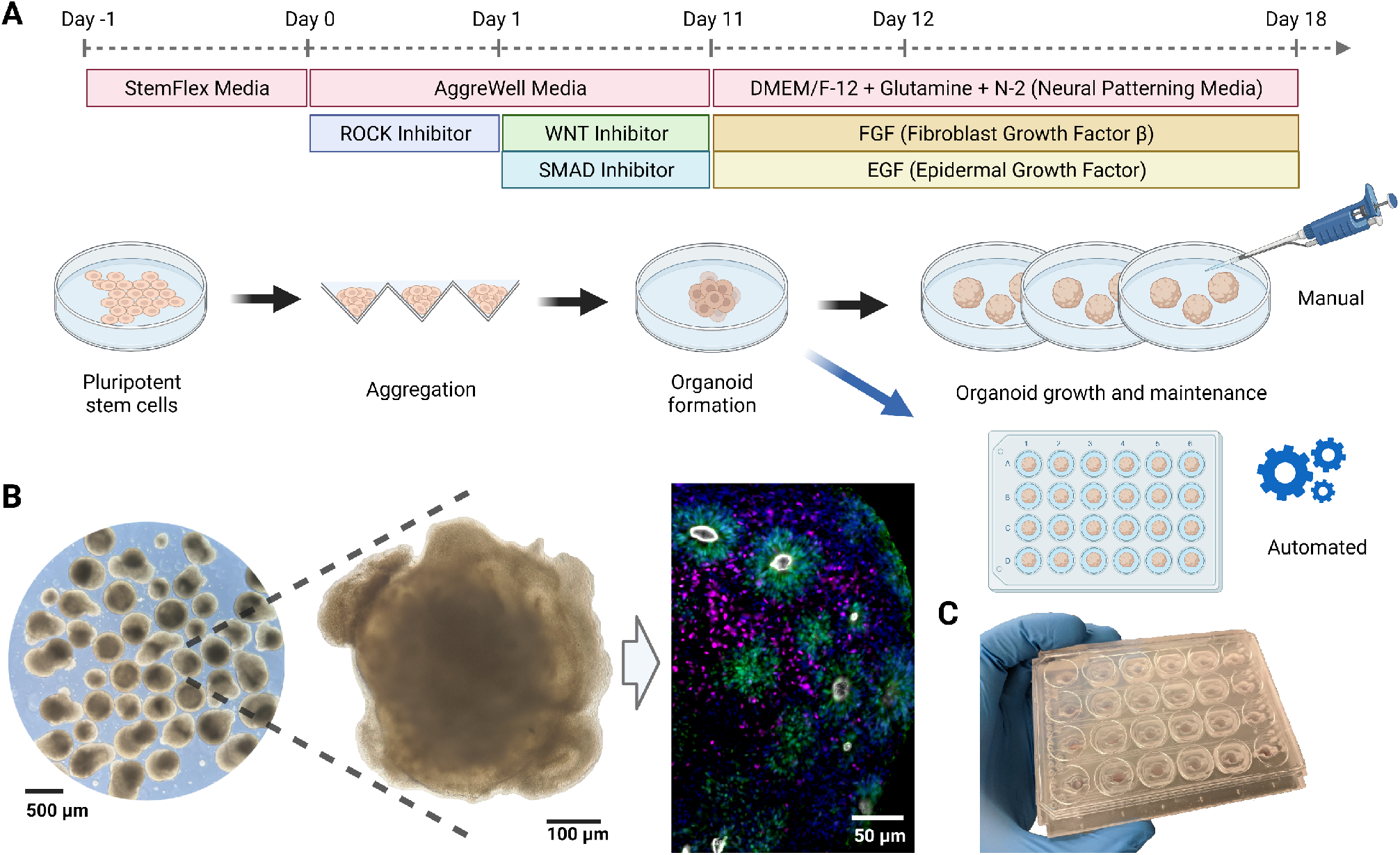
Overview of the human cerebral organoid generation protocol. **(A)** Human pluripotent stem cells are expanded in traditional 2D culture, dissociated, aggregated into microwells, and matured into 3D organoid cultures using defined media conditions to promote cerebral cortex tissue differentiation. In this study, on day 12 post-aggregation, organoids were either kept in suspension and maintained manually (black arrow) or transferred to individual wells of a microfluidic chip and maintained in automation (blue arrow). **(B)** Images of cerebral organoid cultures. Bright-field images at low (left) and high (center) magnification under standard culture conditions show organoid morphology and heterogeneity. Immunofluorescence stains on week 5 for PAX6 (green, radial glia progenitor cells), CTIP2 (BCL11B) (magenta, excitatory projection neurons), ZO-1 (TJP1) (white, tight junction proteins on radial glia endfeet, apical surface of the neural tube), show characteristic ventricular zone-like rosette structures with radial glia surrounded by neurons. Nuclei stained with DAPI (blue). **(C)** Image of the PDMS microfluidic chip. The custom cell culture chip, modeled after a standard 24-well plate, houses organoids for automated experiments.

In conventional cell culture, dispensing, moving, and removing liquid are the necessary actions required in all protocols. Therefore, liquid handling and proper storage of liquid reagents are key features to automate this process. Current liquid handling technologies are broadly classified into two categories: continuous-flow and digital microfluidics^21^. Continuous-flow systems rely on steady-state liquid flow generated by pressure, mechanical, or electrokinetic pumps. With constant flow, these systems deliver a high control of velocity and homogeneity; however, they are less flexible for complex fluidic manipulations and challenging to scale. Closed-channel systems derived from continuous flow are inherently difficult to access, require managing trapped gasses, and do not scale well because the parameters that govern flow at any single location depend on the entire system’s properties.

In contrast, droplet-based, or segmented-flow microfluidics control discrete volumes, which is convenient for droplet manipulation^22^, mixing, encapsulation, sorting, sensing, and high throughput experiments^23^. In combination with valves, droplet-based microfluidics can perform most logical operations upon droplets for sorting, merging, and dividing. Polydimethylsiloxane (PDMS) is the most common material to fabricate microfluidic devices due to its low cost, versatility in prototyping, and high gas permeability^24^. Droplet-based microfluidics has created many opportunities for control that are either impossible or impractical in manual practice, such as scheduling feeding frequency. Within both technologies, there are varieties of *contact* and *non-contact*^25^ approaches. Contact liquid handlers require a tip containing liquid to meet the substrate, similar to pipetting but with greater precision and control. Non-contact liquid handlers manipulate fluid pressures and velocities to control flow. Both technologies have advanced to ultra-low volumes (nano-liter) and ultra-low flow rates (micro-liter per hour).

Organoids are commonly grown in batches with several organoids suspended in single wells of a plate. The shared conditions in a single well support batch consistency; however, increasing the number of variables in an experiment, such as one with multiple genotypes, throughput becomes challenging. Furthermore, multi-well plates are not well suited for controlling dynamic conditions nor generating concentration gradients in the media. In contrast, microfluidic chips as organoid culture chambers enable precise control of environmental conditions with high spatiotemporal resolution^26,27^. Serial dilutions can achieve automated gradients, and robotic volumetric flow can efficiently manage high throughput experiments. The choice of device for laboratory automation should consider the use, flexibility, and cost. Considering the microenvironment of the cell culture, a non-contact system similar to the one developed here offers key advantages to maintaining homeostasis of nutrient concentrations, reducing the build-up of metabolic byproducts, and generating shear forces more similar to *in vivo* conditions.

Here we developed a PDMS-based, non-contact microfluidic platform with segmented feeding to automate prolonged cell culture. The platform is configurable to many organoid protocols and gradient assays. The platform interface enables integration with other research instruments such as an in-incubator imaging system via the Internet of Things. We evaluated this platform by testing its ability to support human cerebral cortex organoid growth and development.

## Results

### The *Autoculture* Platform

#### System Design

We developed an automated, microfluidic cell culture platform to optimize 3D organoid growth, the *“Autoculture”* platform. The system consists of six linked modules (Figure 2): (1) Refrigerator with reagent reservoirs (e.g., fresh media), (2) Syringe pump, distribution valves, and control interface, (3) conditioned media collection reservoirs in cold storage, (4) a microfluidic serial bus interfacing with a cell culture incubator, and (5) a multiplexed microfluidic organoid chip that (6) immobilizes organoids within their micro-environment vessel (1 of 24 wells). To feed the organoids, each individual well is serviced by a fluidic controller; the controller removes spent media via aspiration to a collector in cold storage and then replenishes the vessel with fresh media at programmable intervals.

**Figure 2.**
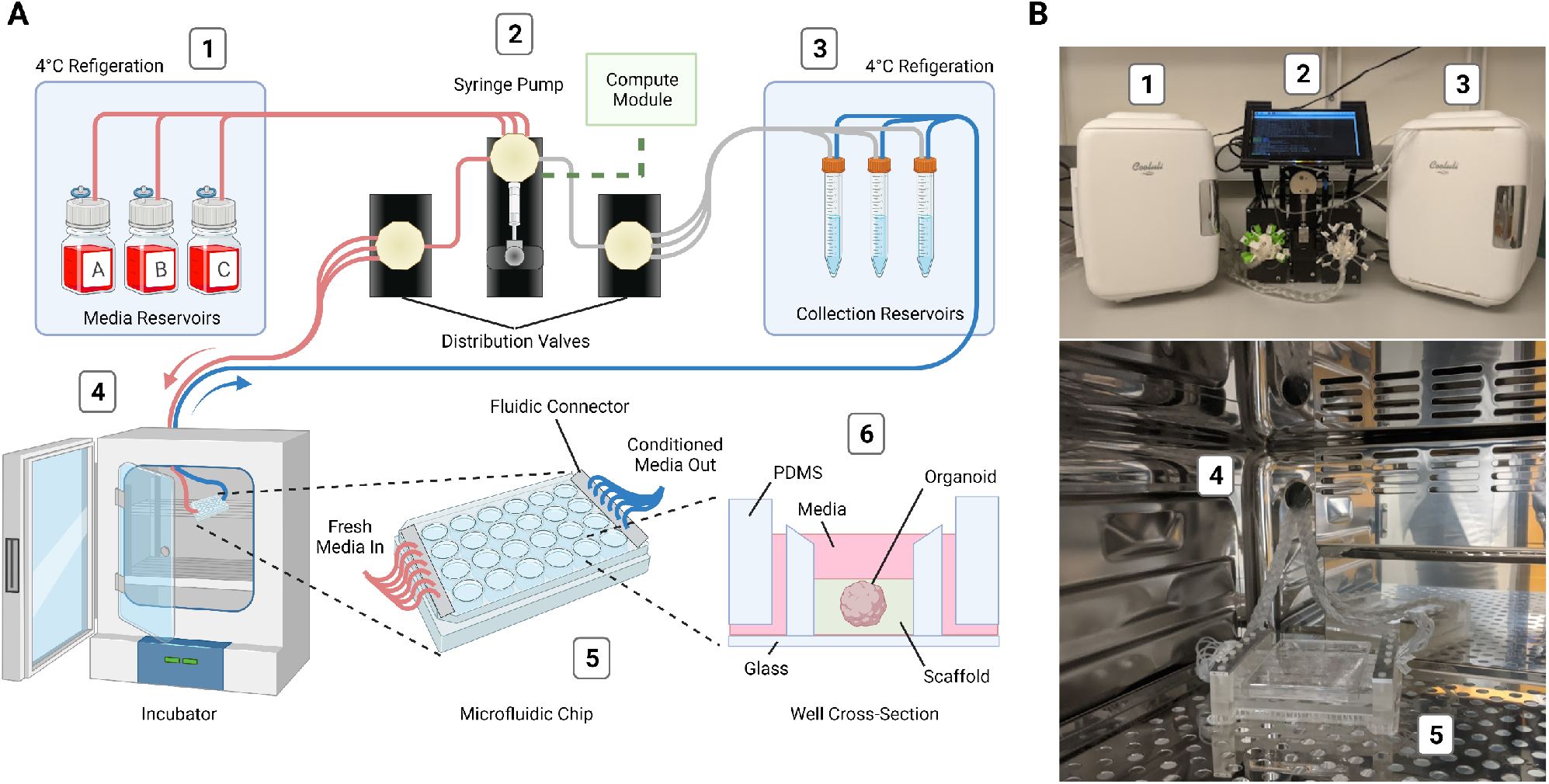
Design and implementation of the automated, microfluidic culture platform. **(A)** Illustration of the automated, microfluidic organoid culture platform, the *Autoculture*. **(B)** Front view images of the *Autoculture*. (1) Refrigerator with reagent reservoirs. (2) Syringe pump, distribution valves, and control interface. (3) Refrigerator with conditioned media collection reservoirs. These components reside on a lab bench directly above the cell culture incubator. (4) Microfluidic tubes enter through an incubator port and connect to the (5) microfluidic well plate chip inside the incubator. (6) Cross-sectional diagram of a single well containing an organoid culture.

Each of the 24 wells of this system is a separate, isolated experiment with a dedicated inlet tube, outlet tube, and collection reservoir. Each well’s feeding schedule is fully customizable in rate and media to increase the flexibility of experimentation. A range of 5-1000 µL aliquots from 3 media/reagent reservoirs can be scheduled to any well. Media/reagent reservoirs may also be used in combination. By design, each well forms a fluidic circuit that maintains isolation from the other circuits on the plate (See Methods). A full 24-well plate is serviced in 72 seconds, and the time between fluid injections may be any length beyond (for instance, every hour, twice a day, every other day, etc.). Conditioned media may be retrieved for molecular analysis without disrupting the culture at any point during or after the experiment. After the experiment, the organoids are retrieved for molecular analysis.

With these control parameters, entire plates may carry out the same workflow to generate consistent batches of organoids. One can also titrate a reagent with an incremental gradient in concentration from well 1 to 24. In addition, one can run multiple protocols/feeding schedules across the plate and change the media components or feeding schedules at various time points throughout an experiment.

#### Internet of Things Connectivity

The Amazon Web Services (AWS) Internet of Things (IoT) provides a convenient control framework for multiple devices to communicate information and initiate actions. With a communication protocol such as Message Queuing Telemetry Transport (MQTT), systems can manage synchronized, coordinated efforts of control and data acquisition in real-time over the internet^28^. System-to-system communication occurs over a local area network (LAN) while researchers can make on-demand calls for data or actions remotely. As such, the *Autoculture* software architecture has control from the AWS Cloud to design, start, pause, edit, and stop experiments on demand (Figure 3).

**Figure 3.**
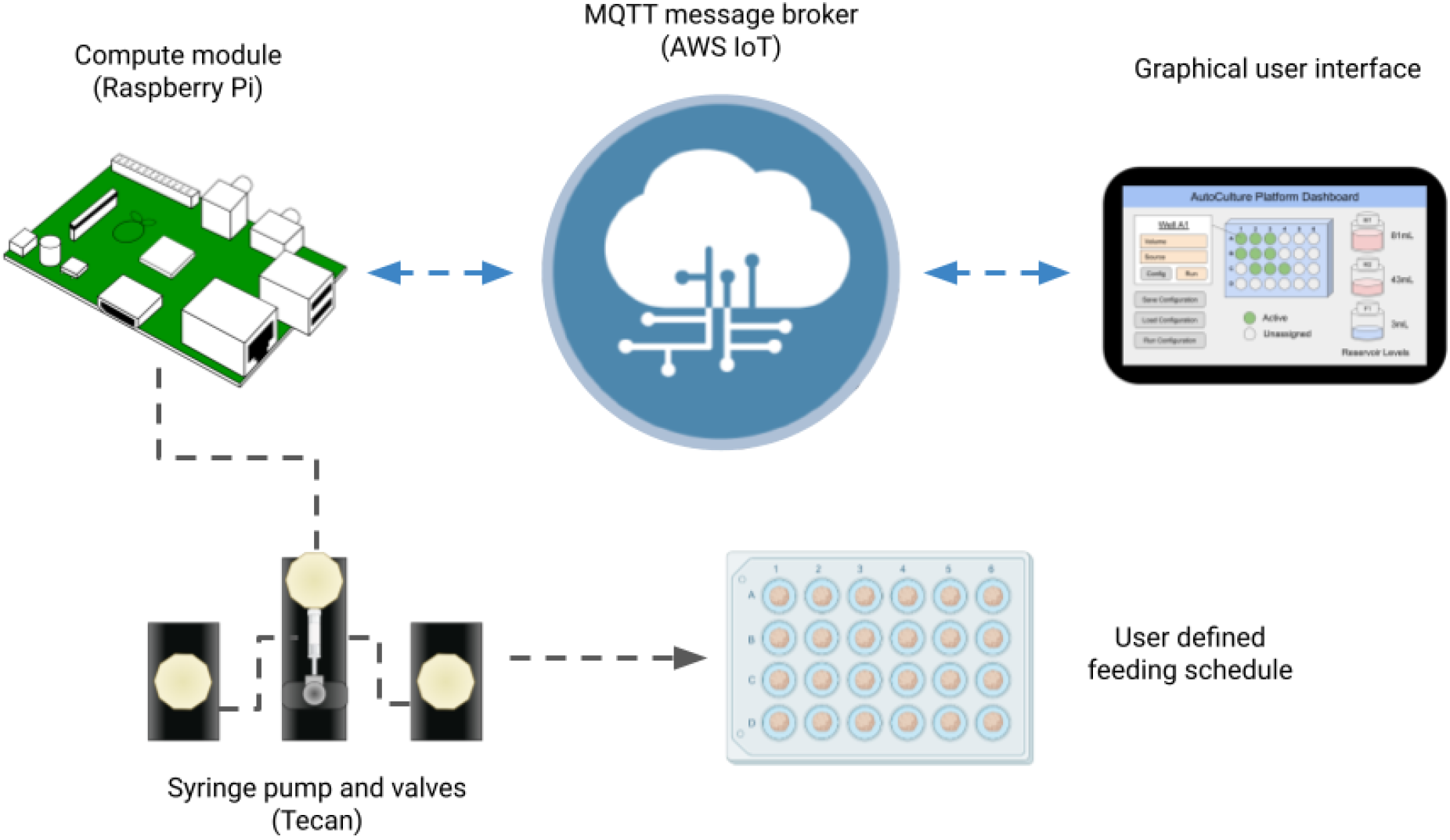
IoT Cloud integration: a graphical user interface hosted on the internet relays messages to the *Autoculture* platform via MQTT to start, monitor and end experiments.

The Raspberry Pi compute module was used to run experiments locally and accept commands relayed over MQTT. The application program interface (API) used to send serial commands to the Tecan pump and valves was adapted from an open-source Python package^29^. Protocols to carry out experiments were developed by timed sequences of unit actions to the pump and valves. As a user, the parameters and ranges available to construct automated experiments are listed in Table 1. In this architecture, IoT connection provides the accessibility to operate experiments remotely while schedules operated locally on-device provide the security of operation in the case of unstable internet connection.

**Table 1.** Experiments run on the *Autoculture* system are configurable in any combination of the listed parameters and ranges.

| Parameter | Range |
| --- | --- |
| Well | 1-24 |
| Media reservoir source | 1-3 |
| Volume in | 5-1000 $\mu\text{L}$ |
| Volume out | 5-1000 $\mu\text{L}$ |
| Maximum feed frequency | 72 sec |

### Microfluidic Organoid Chip

#### Consumable Design

Figure 4 describes the manufacturing and assembly process of the PDMS microfluidic chip. The optically-transparent glass-PDMS microfluidic chip has the footprint of a 24-well plate (85.5 mm x 127.6 mm) to integrate with laboratory tools such as microscopes, plate readers, and robotic dispensers. The 24 isolated wells are addressable via 2 mm square channels at the glass-PDMS interface. For convenient accessibility, all inlets are located in a row on the edge of the chip’s face and all outlets are located in a row on the opposing edge. The open-loop design here is observed with wells that are open to the air. This way, bubbles accumulated in the system are exhausted, there is free gas exchange with the incubator environment, and organoids are easily accessed during chip loading and the experimental conclusion. Each well traps 120 µL that can be used as a micro-environment, including the use of extracellular scaffolds.

**Figure 4.**
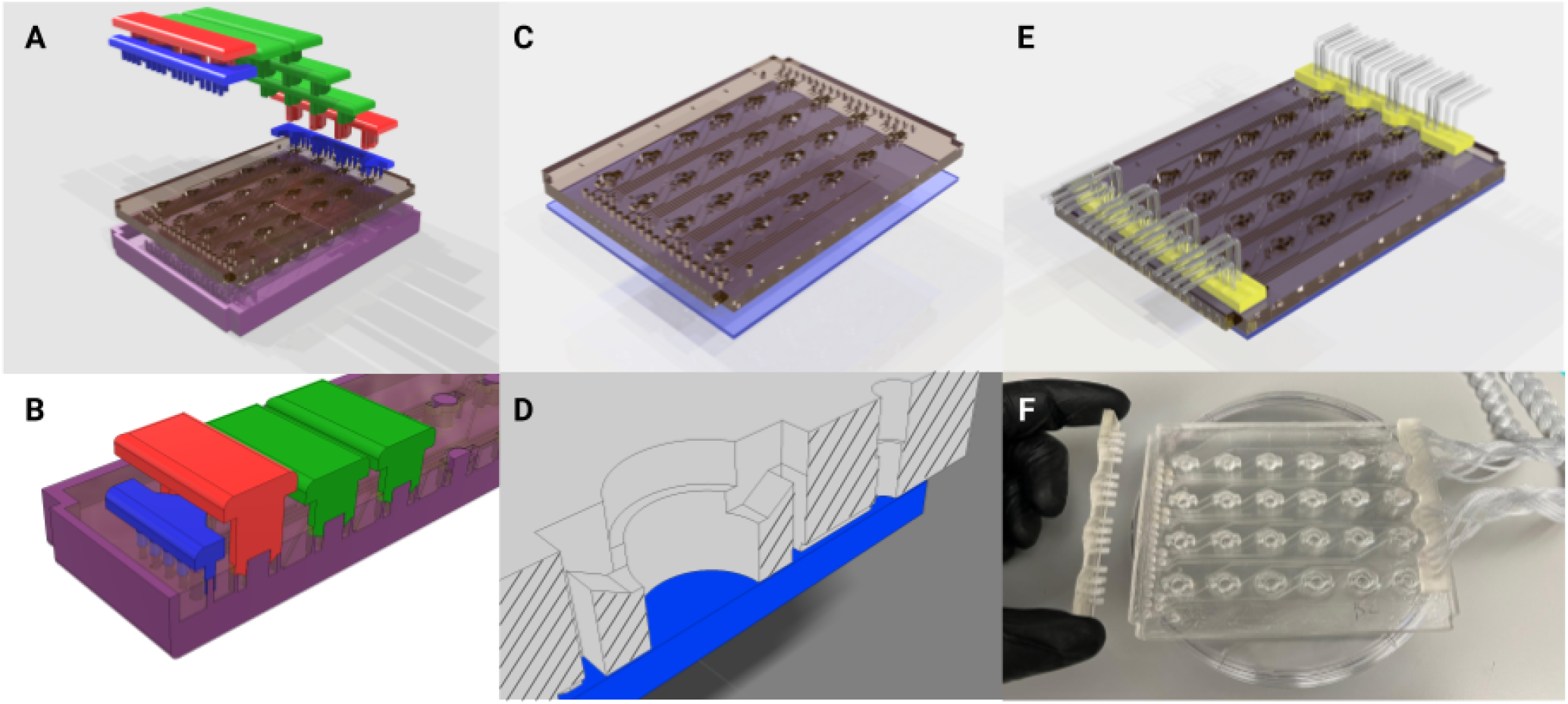
Fabrication of the PDMS microfluidic chip. **(A)** Graphical rendering of the interlocking mold pattern for the PDMS substrate in the microfluidic chip assembly. **(B)** Interlocking mounts (blue, red, and green) affix to the base mold (purple) and define microfluidic geometries upon the poured PDMS that are retained as the substrate cures. **(C)** The PDMS substrate is removed from the mold and bonded to glass. **(D)** A cross-sectional rendering of the chip. Fluid enters from microfluidic inlets on the surface and follows channels sealed by glass on the bottom to wells with open access from the top. **(E)** A 3D-printed fluidic interface plate (yellow) connects 24 fluidic microtube lines with the inlets/outlets of the microfluidic chip. **(F)** Microfluidic chip (center) with an example of the fluidic interface plate (left) and fully installed fluidic interface plate (right).

Each 3D-printed, fluidic interface plate simultaneously connects 24 inlet tubes to the inlets/outlets of the microfluidic chip. Three bores seal the Tygon tubing about the rigid projections. The collection of 24 lines mates with the molded holes in the PDMS, creating a bore seal with all tubes.

#### Fluid Dynamics Simulation

On day 12 of development, cerebral organoid cultures were transferred from orbital shakers (Figure 5A) to automated microfluidic wells (Figure 5B). The orbital shaker maintains a quasi-steady velocity regime of approximately 0.12 m/s^30–32^. We performed a Computational Fluid Dynamics simulation (CFD) using Comsol Multiphysics^33^ (Figure 5C) representing the first 2.2 seconds of media injecting into the well. Fluid enters at the inlet, 5 mm above the glass floor, and enters the well via a sloped wall. The simulation begins with a media liquid domain filled to the level of the inlet/outlet ports with an air domain existing above. In feeding, media enters the cavity and generates a wave that progresses from the inlet to the outlet. The organoid, modeled as a sphere, fixed 5 mm below the air-media interface, does not observe most of the turbulence generated. The lower (glass) substrate experiences <5% of the maximum velocity during feeding. The velocities in this regime are instantaneous (as opposed to quasi-steady-state) and two orders of magnitude less than that of orbital shakers, inducing lower velocities and less shear stress upon the culture. One of the advantages of this system is that researchers can tailor the media flow rate and delivery schedule to achieve optimal diffusion of the nutrients with minimal disruption to the organoid.

**Figure 5.**
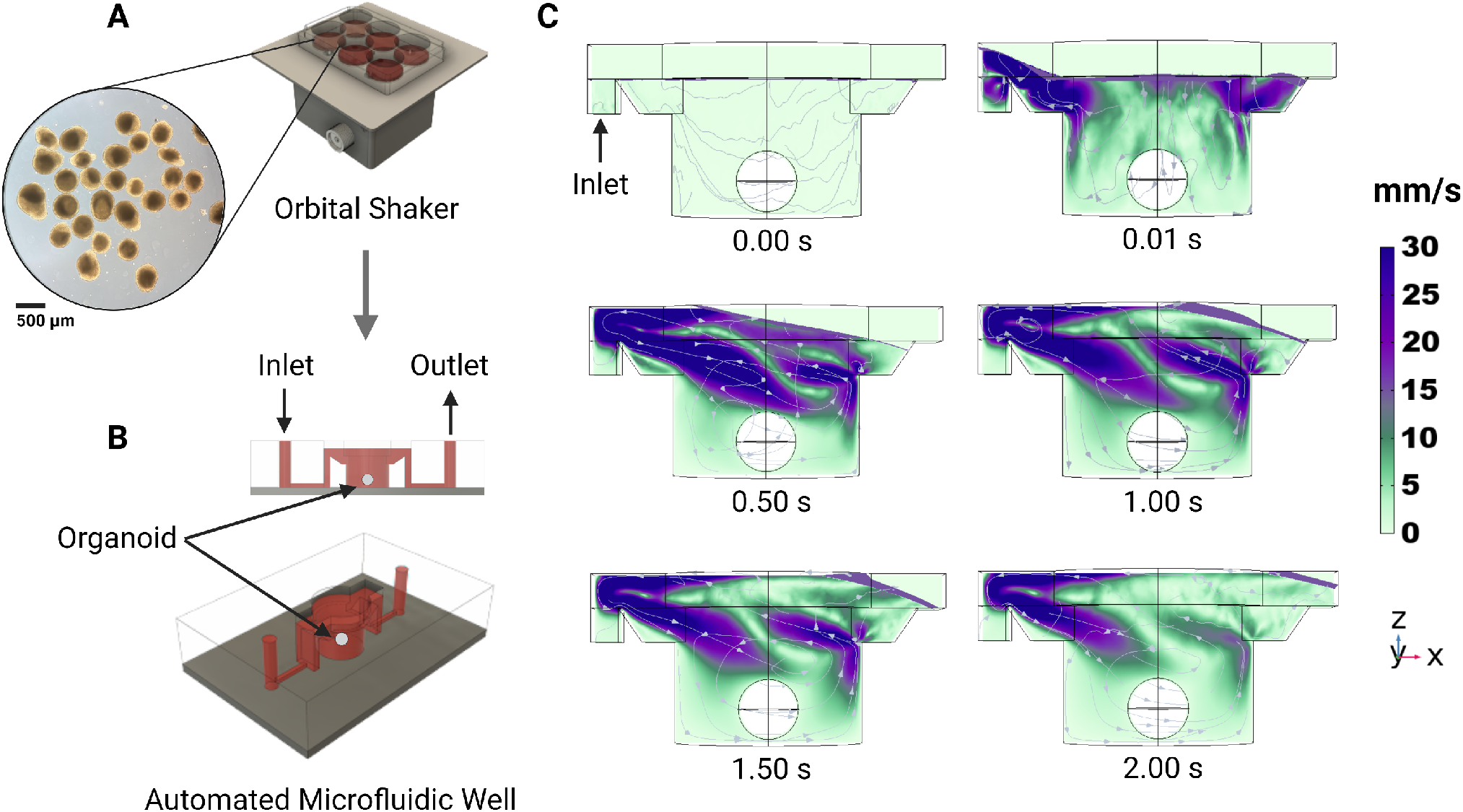
Computational fluid dynamics simulated flow in the individual tissue culture wells using the automated system. **(A)** Organoid cultures are expanded on well plates rotated on an orbital shaker. **(B)** After 12 days, organoids are transplanted into the automated microfluidic well. **(C)** One well of the proposed automated system with constant fluid injection over 2 seconds with velocity streamlines.

### System Validation

#### Culture of Cerebral Organoids

The growth of cerebral organoids on *Autoculture* was compared to that of orbital shaker conditions in an 18-day experiment. Human pluripotent stem cells were aggregated to form organoids and maintained under standard conditions for the first 12 days during neural induction (see *Methods*). The batch was split, and 12 organoids were loaded onto an *Autoculture* microfluidic chip while the remainder were maintained in a 6-well plate on an orbital shaker as controls. Each automated organoid was fed 70 µL every hour for six days, while the controls were fed 2 mL every other day. In automation, the well plate does not need to be periodically removed from the incubator for feeding; therefore, this system is well-suited for longitudinal monitoring of organoid development. In this study, organoids in the *Autoculture* well plate were monitored once per hour using a bright-field imaging in-incubator platform^34,35^.

#### Live Imaging

Figure 6A shows the automated, microfluidic culture plate inside an incubator placed on a remote-controlled, IoT-enabled, multi-well automated imaging system. The imaging system was designed to monitor biological experiments in a 24-well plate format, using one dedicated camera for each well. To account for the three-dimensional development of the organoids during the entire experiment, we captured bursts of images, sweeping through the range of focal distances covering the entire three-dimensional tissue. A computer vision algorithm was used to detect the features in focus at each focal plane, generate a composite image maximizing the in-focus features in the entire organoid, and compute the projected area. This process is described in previous work^35^. Figure 6B shows 12 cerebral cortex organoid cultures (day 12), loaded in individual wells of the microfluidic chip and fed in parallel on the *Autoculture* platform for the experiment. Figure 6C and D show the growth of “Culture 4” over six successive days. Robust organoid growth was observed for the organoids in the *Autoculture* wells and was consistent with the size increase observed for the organoids grown under control conditions. Compared to controls, automated organoids did develop a less-dense perimeter, suggesting that the reduction in velocities and shear forces may accommodate growth and migration that would otherwise be cleaved.

**Figure 6.**
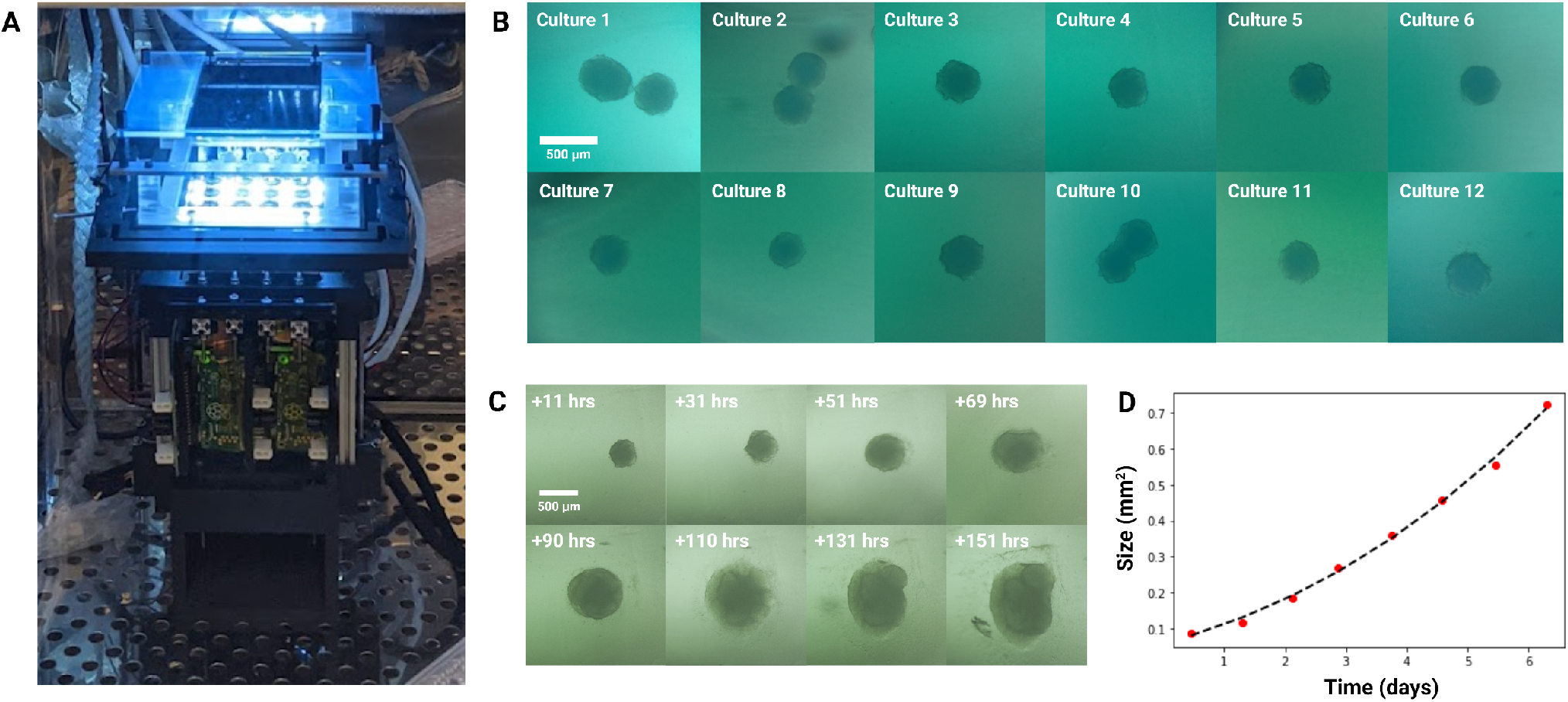
Longitudinal monitoring of organoid development. **(A)** The *Autoculture* microfluidic chip sits on a remote-controlled, IoT-enabled, 24-well automated imaging system. **(B)** Bright-field images of twelve individual 12-day-old cerebral cortex cultures at day 1 of automated feeding. **(C)** Longitudinal imaging of “Culture 4” during the experiment. **(D)** Projected area expansion of “Culture 4” during the experiment. This was obtained using a computer vision algorithm^34^.

#### Growth Analysis

On day 18, the cultures were harvested and analyzed by bulk RNA sequencing (RNA-seq) and immunohistochemistry to evaluate the cell types generated in each condition and the overall health of the cell cultures. The transcriptomes of 7 “Automated” and 4 “Suspension” organoids were compared. Gene expression of cell-type markers for neuralepithelia (SOX2), radial glia neural stem cells (SOX2, HES5, PAX6, HOPX), intermediate progenitors (EOMES), and immature neurons (NeuroD1, RELN) did not show consistent differences between the *Autoculture* and control samples suggesting that overall differentiation fidelity was not affected by using the *Autoculture* system. Consistent with this, we saw a robust expression of the neural progenitor protein markers, SOX2 and Nestin by immunohistochemistry in sections of organoids grown under standard or *Autoculture* conditions (Figure 7B).

**Figure 7.**
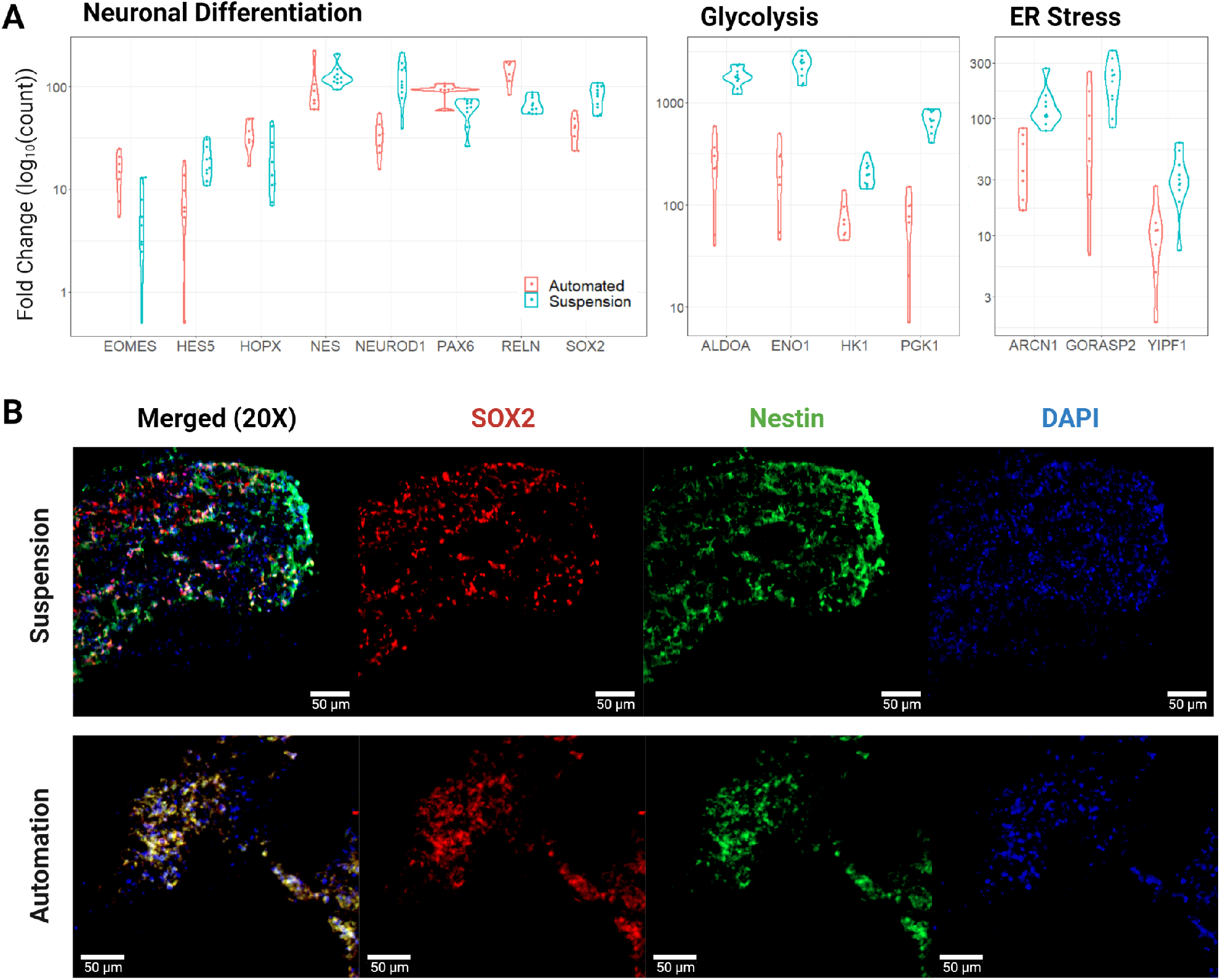
Transcription and immunofluorescent imaging results. (A) Pairwise comparisons of expression for select genes associated with neural differentiation, glycolysis, and ER stress. Results were statistically significant with an adjusted p-value <= 0.05, except for HOPX and NES of neural differentiation. The “Automated” data represent 7 biological replicates and the “Suspension” data represent 4 biological replicates with 6 technical replicates **(B)** Immunofluorescent stains for SOX2, Nestin, and DAPI of Suspension and Automated organoid sections show congruent progenitor markers.

Contrasting automated and control organoids, we observed a significant difference in the expression of genes associated with cell culture stress. Studies have shown that glycolysis and endoplasmic reticulum stress (ER stress) pathways are upregulated in organoids relative to *in vivo* tissue samples, and these correlate with impaired cell subtype specification and maturation^17,36^. In this study, canonical glycolysis was the top deferential pathway with the vast majority of the significant genes consistently downregulated (see Supplementary Tables 2 and 3). Genes that are notably upregulated in cerebral organoids showed reduced levels in the automation condition: ALDOA, ENO1, HK1, and PGK1 had fold changes of -6.9, -10.4, -2.8, and -9.3, respectively (Figure 7A). Additionally, markers of ER stress: ARCN1, GORASP2, and YIPF1 were reduced by fold changes of -2.8, -2.2, and -3.0, respectively. See Supplementary Table 4 for more deferentially expressed genes of ER stress.

## Discussion

The increasing demands for long-term experiments, reproducibility, parallelization, and longitudinal analysis drive cell culture toward automation. This study showcases an automated, microfluidic solution for the growth and maintenance of organoids capable of existing in conjunction with other control and sensing devices over the Internet of Things, magnifying the ability to capitalize on the precision robotics for automated experimentation. Combining this platform with the imaging platform shown here (Figure 6) provides a stationary environment that uniquely enables the live study of individual organoids over time.

The use of orbital shakers has been widely adopted in the field and yields benefits to nutrient diffusion and dissolved gas homogeneity. However, the fluid velocities and shear forces present do not resemble the embryonic environment. Microfluidics such as in the system shown here could be used to tune fluid velocities, shear forces, and pressure gradients while providing the same benefits to nutrient and dissolved gas concentration through rapid feeding regimens. Our system enables an environment of low velocities. The CFD simulation shown here (Figure 5) represents the conditions present in the validation experiment which preferentially selected for low velocities; however, greater rates of profusion would yield higher velocity gradients that could be used to study the effects on organoid morphology and differentiation.

Reducing stress levels in cerebral organoid models is crucial to achieving physiological relevance. As measured by reduced glycolytic enzyme expression, the environment of the *Autoculture* platform results in reduced-stress organoids compared to traditional suspension culture conditions. Pathways that respond to environmental conditions such as sugar metabolism, hypoxia monitoring, and protein production are interconnected through the integrated stress response pathway. By reducing concentration fluctuations in the cell culture media through automated feeding, the cultures may experience greater homeostasis. Further investigation is needed to understand the potential long-term effects of reducing gene expression of glycolysis and ER stress genes and the critical environmental conditions that underlie the gene expression signature associated with less cell stress we observed. For example, it is unclear whether depletion of essential nutrients, like glucose, or accumulation of cell metabolites, like lactose, is the critical factor leading to the induction of genes in the glycolysis pathway observed under standard organoid culture conditions. However, the *Autoculture* system we developed here provides a platform in which we can systematically explore this question.

## Methods

### System Design and Assembly

#### *Design of the* Autoculture *platform*

Cell culture media was stored in glass bottle reservoirs (Corning) with a multi-port solvent delivery cap (Spex VapLock) and stored at 4°C for the duration of the experiment. Each reservoir delivery cap contained a single 0.030” ID x 0.090” OD Tygon microbore tube (Masterflex), sealed by a two-piece PTFE nut and ferrule threaded adapter (Spex VapLock), extending from the bottom of the reservoir to an inlet port on the 6-port ceramic valve head of the syringe pump (Tecan Cavro Centris, 1.0-mL glass vial). Sterile air is permitted to backfill the reservoir through a 0.22-µm filter (Millipore) affixed to the cap to compensate for syringe pump reagent draws. The same Tygon microbore tubing and PTFE nut and ferrule threaded adapters were used to connect the syringe pump to two parallel 12-port distribution valves (Tecan SmartValve). Each 0.020” ID x 0.060” OD Tygon microbore tube (Masterflex) emanating from the distribution valve connects to a single well of the microfluidic chip. Fluidic isolation between wells is retained from this junction onward. Each 12-port distribution valve services six wells on the microfluidic chip. Systems with two and four distribution valves were constructed. The collection of 2-meter long microbore tubes was bundled into a braid for convenient handling and guided through the rear entry port of a standard cell culture incubator (Panasonic) (Figure 2(4)). At incubator conditions (37°C, 5% CO2, 95% rel. humidity), a single custom, 3D-printed fluidic interface plate mated the set of microbore tubes for reagent delivery to the inlets of the microfluidic chip and a second, identical interface plate mated the set of microbore tubes for reagent aspiration to the outlets.

The microbore tubes for aspiration were guided back out of the incubator and to a set of single-use 15-mL conical tubes (Falcon) for conditioned media collection (Figure 2(3)). Each collection reservoir was capped with a rubber stopper (McMaster) containing two 0.06” drilled holes. For each stopper, the microbore tube sourcing conditioned media from the microfluidic chip was inserted into one hole, and a dry microbore tube for pneumatic operation routes back to the aspiration distribution valve head was inserted into the other hole. The syringe pump was used to generate negative pressure upon each collection reservoir in series to draw conditioned media from the microfluidic chip into the collection reservoir. The separation between the pneumatic tube and conditioned media tube trapped the influx of media in the collection reservoir. All microbore tubes were hermetically sealed to the distribution valve and syringe pump with PTFE nut and ferrule threaded adapters.

The syringe pump and distribution valves were connected using the multi-pump electrical wiring configuration described in the Tecan manual. A compute module (Raspberry Pi 4) relayed serial communications with Tecan OEM Communication Protocol using a GPIO TX/RX to DB9M RS232 serial expansion board for Raspberry Pi (Ableconn). The Raspberry Pi compute module used a 7” touchscreen display to edit and launch protocols. An open-source Python application program interface (API) was used to develop the software required to carry out protocols in automation^29^.

#### PDMS Molding

PDMS-based microfluidics were constructed using an interlocking 3D-printed plastic mold (Figure 4). These were printed with an SLA printer (Formlabs Form 3) with Model V2 resin. The printed molds were post-processed with sonication in isopropanol (IPA) for 20 minutes to remove excess resin, followed by drying in N2. Dry components were cured under UV-light (405nm) for 30 minutes at 60°C. As illustrated in Figure 3A, the mold parts were assembled and filled with PDMS (Sylgard 184, Dow Corning) prepared by mixing PDMS prepolymer and a curing agent (10:1 w/w). After filling the mold, the PDMS was degassed in a vacuum chamber for 1 hour. The PDMS-filled mold is left to cure for 24 hours at 60°C before removal of the PDMS from the mold.

#### Microfluidic Chip Assembly

Borosilicate glass substrates (101.6mm x 127.0mm, McMaster-Carr) were cleaned via sonication in acetone (10 minutes), then isopropyl alcohol (10 minutes), and dried with N2. The glass substrate and molded-PDMS surface were activated with oxygen plasma at 50W for 45 seconds. The glass and PDMS (Figure 4C) were aligned by hand, pressed together, and baked at 100°C on a hot plate for 30 minutes, forming an irreversible seal.

#### Parylene Coating

A 10 µm layer of parylene-C (Specialty Coating Systems) was deposited onto the microfluidic chip to prevent PDMS absorption of small molecules. Two drops of silane A-174 (Sigma-Aldrich) were loaded into the deposition chamber to promote adhesion.

#### 3D Printed Components

The fluidic interface plate (Figure 4F) was 3D printed to interface the 2.2 mm OD microbore tubing (Cole-Palmer) with the PDMS inlet and outlet features. Each connector geometry consisted of 24 cylindrical extrusions with an OD of 2.7 mm and a bore of 2.2 mm. Within each bore were three 0.2 mm long barbs to grip the microbore tubing when inserted. This component was printed with a Formlabs SLA printer (Form 2) with Surgical Guide resin with sonication in isopropanol (IPA) for 20 minutes to remove excess resin, followed by drying in N2. Dry components were then cured under UV light (405 nm) for 30 minutes at 60°C. To ensure biocompatibility, the part was coated with 5 µm of parylene-C (Specialty Coating Systems). Two drops of silane A-174 (Sigma-Aldrich) were also loaded in the deposition chamber with the device to promote adhesion.

### Organoid Cell Culture

#### Sterilization

Sterilization of the syringe pump, valve heads, tubing, fluidic interface plates, and collection tube caps was carried out per supplier recommendation (Tecan). A 10-minute wash with 70% ethanol sourced from an autoclaved glass reservoir was pushed through the platform to the collection reservoirs via the syringe pump. Following 70% ethanol, a drying cycle of sterile air sourced through a 0.22 µm filter (Millipore) was applied for 10 minutes. Ten subsequent cycles of deionized, nucleus-free water and drying were carried out under the same parameters. The microfluidic chip and media reservoirs were autoclaved at 121°C for 45 minutes and dried for 15 minutes immediately prior to use. All components were transferred via sealed autoclavable bags to a biosafety cabinet in tissue culture for organoid and media loading. Pre-prepared, supplemented media was transferred to each media reservoir and capped (VapLock) and then stored in refrigeration for the duration of the experiment.

#### hESC line maintenance

The human embryonic stem cell line H9 (WiCell, authenticated at source) was grown on recombinant human vitronectin (Thermo) coated cell culture dishes in StemFlex Medium (Gibco). Subculturing was performed by incubating plates with 0.5 mM EDTA for 5 minutes and then resuspended in culture medium to be transferred to new coated plates.

#### Cerebral organoid differentiation and protocol

To generate cerebral organoids, adherent cultures were dissociated into single cells using Accutase Cell Dissociation Reagent (Gibco) and then aggregated in AggreWell 800 24-well plates (STEMCELL Technologies) at a density of 3,000,000 cells per well with 2mL of AggreWell Medium (STEMCELL Technologies) supplemented with Rho Kinase Inhibitor (Y-27632, 10 µM, Tocris, 1254) (day 0). The following day (day 1), 1mL of the AggreWell medium was manually replaced with supplemented medium containing WNT inhibitor (IWR1-ε, 3 µM, Cayman Chemical, 13659, days 1-10) and Nodal/Activin inhibitor (SB431542, Tocris, 1614, 5 µM, days 1-10). On day 2, aggregates were transferred onto a 37 µm filter (STEMCELL Technologies) by carefully aspirating with a p1000 wide-bore pipette out of the AggreWell plate. The organoids were transferred into ultra-low adhesion 6-well plates (Corning) by inversion and rinsing of the filters with fresh AggreWell medium. Media was changed on days 3, 4, 5, 6, 8, and 10 by manually replacing 2 mL of conditioned media with fresh media. On day 11 and onward, the medium was changed to Neuronal Differentiation Medium containing Eagle Medium: Nutrient Mixture F-12 with GlutaMAX supplement (DMEM/F12, Thermo Fisher Scientific, 10565018), 1X N-2 Supplement (Thermo Fisher Scientific, 17502048), 1X Chemically Defined Lipid Concentrate (Thermo Fisher Scientific, 11905031) and 100 U/mL Penicillin/Streptomycin supplemented with 0.1% recombinant human Fibroblast Growth Factor b (Alamone F-170) and 0.1% recombinant human Epidermal Growth Factor (RD systems 236-EG).

Control-group “Suspension” organoids remained suspended in 6-well plates and were maintained with 2 mL media changes every other day for the remainder of the culture. Experimental-group “Automated” organoids were loaded onto the microfluidic chip and experienced media changes of 70 µL once every hour for the remainder of the culture.

#### Microfluidic chip loading

On day 12 of cerebral organoid differentiation, the microfluidic chip was prepared by pipetting 50 µL of chilled (approximately 0°C) Matrigel hESC Qualif Matrix (BD 354277) into each well. Immediately following Matrigel, single cerebral organoids were transferred via p1000 wide-bore pipette with 70 µL of native conditioned media to each well and positioned to center-well for imaging. The chip was covered with a 24-well plate lid and incubated at 37°C for 15 minutes to set the Matrigel. Each well was filled with an additional 70 µL of fresh media and connected to fluidic interface plates (Figure 4F) routed into the incubator through a rear access port. The fluidic interface plates were pressure fitted into the microfluidic chip by hand, and the chip was positioned on the imaging platform (if applicable).

### Analysis

#### Sequencing library preparation

Smart-seq2 protocol^37^ was used to generate full-length cDNA sequencing libraries from whole organoid mRNA. Briefly, whole organoids were lysed using lysis buffer to render cell lysate containing polyadenylated mRNAs that were reverse transcribed with Superscript III (ThermoFisher Scientific) using an oligoDT primer (/5Me-isodC/AAGCAGTGGTATCAACGCAGA GTACTTTTTTTTTTTTTTTTTTTTTTTTTTTTTTVN) and template switching was performed with a template switch oligo (AAGCAGTGGTATCAACGCAGAGTACATrGrGrG). The oligoDT primer and template switch oligo sequences served as primer sites for downstream cDNA amplification (AAGCAGTGGTATCAACGCAGAGT). cDNA was quantified using a Qubit 3.0 DNA high sensitivity fluorometric assay, and quality was assessed using a bioanalyzer DNA high sensitivity kit (Agilent). Nextera HT transposase (Illumina) was used to convert 1 ng of cDNA into barcoded sequencing libraries.

#### Transcriptome analysis

Paired-end reads were sequenced at 75×75 bp on an Illumina NextSeq 550, and further depth was sequenced at 50×50 bp on an Illumina NovaSeq 6000 to an average read depth of 65 million paired reads per sample. Samples were demultiplexed using Illumina i5 and i7 barcodes, and higher depth samples were sub-sampled to 100M using SAMtools^38^. Trimmed reads were combined and aligned to the human genome (hg38 UCSC assembly) with STAR alignment^39^ (Gencode v37) using the toil-rnaseq pipeline^40^. STAR parameters came from ENCODE’s DCC pipeline^41^. Differential gene expression was performed using the DESeq2^42^ package in RStudio. Gene set enrichment analysis was performed using g:Profiler^43^.

#### Immunostaining

Cerebral organoids were collected, fixed in 4% paraformaldehyde (ThermoFisher Scientific 28908), washed with 1X PBS and submerged in a 30% sucrose (Millipore Sigma #S8501) in PBS solution until saturated. Samples were embedded in cryomolds (Sakura - Tissue-Tek Cryomold) containing tissue freezing medium (General Data, TFM-C), frozen and stored at -80°C. Organoids were sectioned with a cryostat (Leica Biosystems #CM3050) at 18 µM onto glass slides. Organoids sections were washed three times for ten minutes in 1X PBS prior to a two-hour incubation in 10% BSA in PBS blocking solution (ThermoFisher Scientific BP1605-100). The sections were then incubated in primary antibodies diluted in blocking solution overnight at 4°C. The next day, sections were washed three times with 1X PBS for thirty minutes. They were then incubated in a solution of secondary antibodies diluted in a blocking solution at room temperature for two hours. The sections were washed an additional three times in 1X PBS for thirty minutes.

Primary antibodies used were: rabbit anti SOX2 (ab97959, 1:250 dilution), chicken anti Nestin (ab134017, 1:250 dilution), and DAPI (Sigma D9542-10mg). Secondary antibodies used were: goat anti-rabbit Alexa Fluor 594 (ab150080, 1:250 dilution) and goat anti-chicken Alexa Fluor 488 (ab150169, 1:250 dilution). Imaging was done using the Zeiss Axioimager Z2 Widefield Microscope at the UC Santa Cruz Institute for the Biology of Stem Cells (RRID:SCR021135) and the Zen Pro software. Images were processed using ImageJ.

#### Computational fluid dynamics

The fluid dynamics of filling and draining the wells were predicted using a commercial Computational Fluid Dynamics software COMSOL® Multiphysics 5.5 (Stockholm, Sweden). Figure 5B shows the first 2 seconds of the filling cycle (70*µL* of media delivered with an average velocity of 9.85 × 10^4^m/s). The well is 5mm deep and has a diameter of 5.6mm. In this simulation, the media properties were 997*kg/m*^3^ density and 6.92 × 10^3^kg/m.s viscosity. The simulation predicts the phase boundary (between liquid and air) as a free surface^44^. The solution domain consists of a rigid wall (the well) with “non-slip” boundary conditions and the top surface (the air-media interface) open to the incubator with “slip” boundary conditions. The organoid was simulated as a phantom sphere geometry with a 1.8mm. Atmospheric conditions were set to a pressure of 1atm, a temperature of 37°C, and a gas composition of 5% carbon dioxide, 17% oxygen, and 78% nitrogen. We visualized the velocity field on the central vertical cross-section of the well using stream arrow lines. The simulation used a total of 519, 830 tetrahedral elements.

## Data availability

Polyadenylated RNA sequencing data have been deposited in GSE207894. Additional differential expression data that sourced Figure 7A is available in Supplementary Tables 1-4.

## Acknowledgements

This work was supported by the Schmidt Futures Foundation SF 857 and the National Human Genome Research Institute under Award number 1RM1HG011543 (D.H., S.R.S and M.T.), the National Institute of Mental Health of the National Institutes of Health under Award Number R01MH120295 (S.R.S.) Defense Advanced Research Projects Agency (DARPA), Army Research Office, under Cooperative Agreement No. W911NF-18-2-0104, and the Department of Interior, Award No. D20AC00003. (M.R. and M.T), the National Science Foundation under award number NSF 2034037 (R.E.G., S.R.S and M.T) and NSF 2134955 (M.T., S.R.S and D.H). D.H. is a Howard Hughes Medical Institute Investigator. The authors want to give a special thanks to Pattawong Pansodtee, David Parks, Matt Elliott for discussions and the IBSC Cell Culture Facility (RRID:SCR 021353), and the UCSC Life Sciences Microscopy Center (RRID:SCR 021135) for valuable resources and assitance.

## Author contributions statement

S.T.S, J.S., S.C. and F.A. developed the automated microfluidic cell culture platform. G.L.M. and R.N.H. provided cell cultures. S.T.S and G.L.M. performed the experiments and transcriptome analysis. S.T.M. developed the computational fluid dynamics simulation P.V.B. and V.T.L. performed image acquisition and analysis. L.T. and R.N.H. performed immunohistochemical staining. M.R., D.H., S.R.S., R.E.G. and M.T. supervised the team and secured funding. All authors contributed to the writing of the manuscript.

